# Local Ecological Knowledge enhances our capacity to document potential climate sentinels: a loggerhead sea turtle (*Caretta caretta*) case study

**DOI:** 10.1101/2023.11.01.565199

**Authors:** Michelle María Early-Capistrán, Nicole L. Crane, Larry B. Crowder, Gerardo Garibay-Melo, Jeffrey A. Seminoff, David Johnston

## Abstract

Climate change is inducing rapid transformations in marine ecosystems, with a pronounced effect on top predators like loggerhead sea turtles (*Caretta caretta*). Loggerheads’ responsiveness to temperature fluctuations underscores their significance as a climate sentinel species. We present the first record of a loggerhead sea turtle in Monterey Bay, California, USA, documented through Local Ecological Knowledge (LEK) and community or citizen science (CS), highlighting the pivotal role of these approaches in documenting species occurrences beyond anticipated habitats during climatic anomalies. In rapidly changing conditions, rigorously documented CS and LEK offer a crucial complement to conventional scientific methods, providing high-quality data with extensive coverage— especially for elusive species—and yielding insights into emerging phenomena.

---

Climate change is leading to rapid and unprecedented biological shifts in marine environments (Beaugrand et al., 2019). Marine top predators, including carnivorous sea turtles, are important sentinels of climate change impacts on marine environments given their clear responses to environmental variability (Hazen et al., 2019). Climate change is shifting species distributions, and may cause ecosystems and abiotic drivers to vary faster than time-frames required for scientific field research (Taheri et al., 2021). These difficulties are amplified for highly-migratory endangered species, which are challenging to sample and model, especially outside established habitats, due to their rarity (Jeliazkov et al., 2022). Local Ecological Knowledge (LEK)—place-based knowledge held by specific people about their surrounding environment (Bélisle et al., 2018)—and community or citizen science (CS), data collection by the general public for scientific research—are critical for locating sentinel species in unexpected locations (Tengö et al., 2021; Hanna et al., 2021).

In this context, we present the first record of a loggerhead sea turtle (*Caretta caretta*, henceforth “loggerhead”) in Monterey Bay (MB), California, U.S.A. (36.804358°, −121.786828°), ~1,500 km from the nearest resident population (Figure 1). MB is a marine vertebrate biodiversity hotspot characterised by seasonal upwelling and high productivity (NOAA, 2023a). The loggerhead sighting occurred near the edge of Monterey Submarine Canyon, with maximum depths >2300m (Paull & Caress, 2019), and were registered through LEK and CS. To date, four other hard-shelled sea turtle sightings have been reported in MB (*Chelonia mydas* in 2011 and *Lepidochelys olivacea* in 1969, 2011, and 2023) (See Supplementary Material).

**Figure 1:**
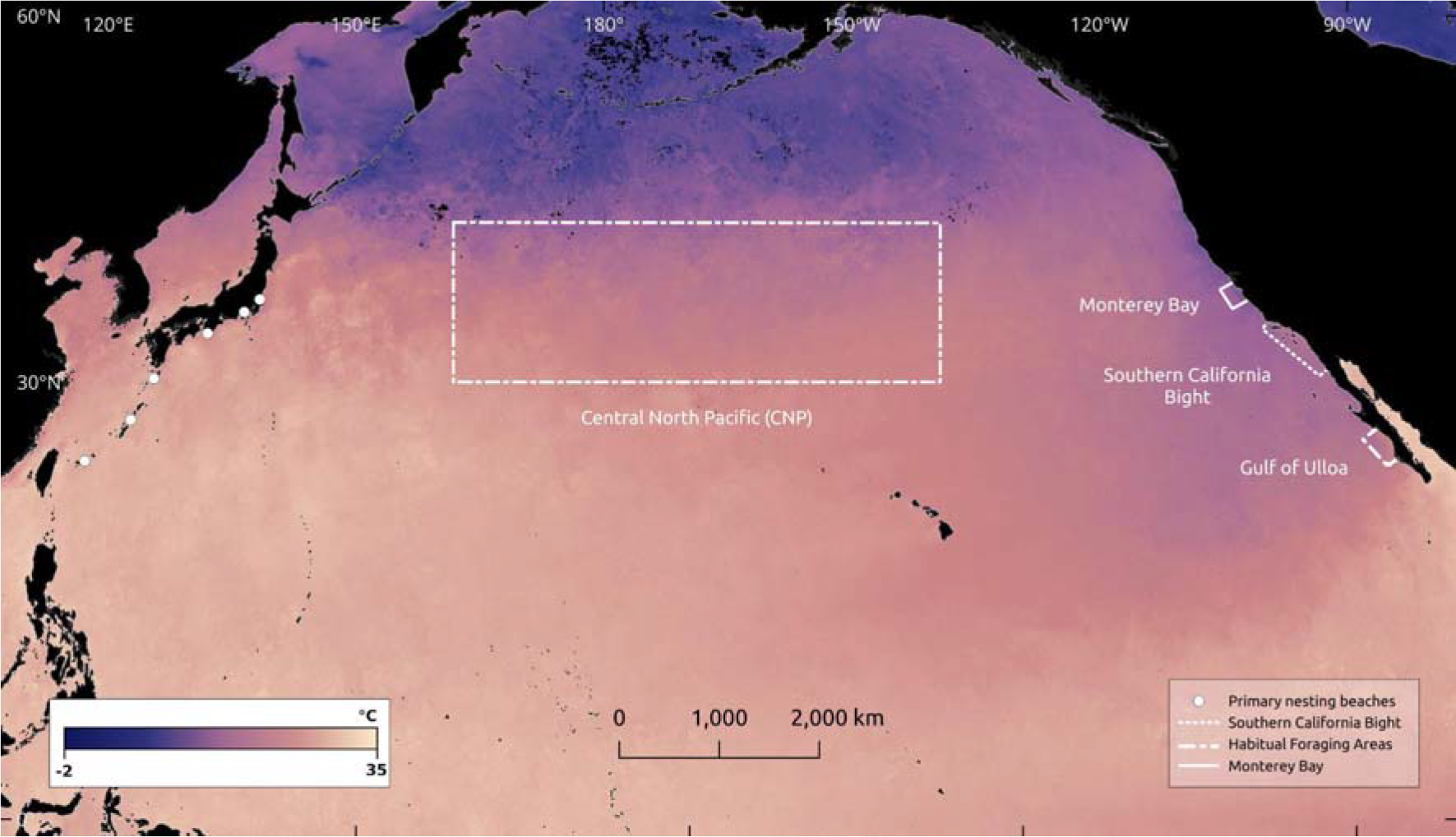
Loggerhead (*Caretta caretta*) nesting beaches, predicted foraging habitats, Monterey Bay and Southern California Bight with July, 2023 Sea Surface Temperature at 0.1° resolution (NASA Earth Observatory, 2023).

Loggerheads are categorised as *Vulnerable* on the IUCN Red List, and their close connection to temperature variability makes them valuable sentinels of temperature anomalies or shifting species range due to climate change (Hazen et al., 2019). North Pacific loggerheads nest exclusively in Japan and forage in highly productive pelagic zones in the Central North Pacific and Baja California Sur, Mexico (Briscoe et al., 2021) (Figure 1). As ectotherms, their distribution and migratory patterns are largely driven by ocean temperature, with important links to annual variations, irregular events (El Niño Southern Oscillation, ENSO), and decadal patterns (Pacific Decadal Oscillation) subject to climate change-driven alteration (Almpanidou et al., 2019; Briscoe et al., 2021).

The loggerhead sighting was reported by co-author D. Johnston, a naturalist with >40 years of experience and ~16,000 hours of observation in MB. M.M. Early-Capistrán and G. Garibay-Melo employed semi-structured interviews and participatory modelling to collaboratively create spatial representations of LEK (Gray et al., 2017). Species identification followed established ethnobiological methods using visual stimulus (Albuquerque et al., 2014), relying on three sea turtle identification guides and composite images of potential species (*C. caretta, L. olivacea*, and *C. mydas*). Leatherback turtles (*Dermochelys coriacea*) were excluded due to their distinct morphology (Benson et al., 2020) (see Supplementary Material). The discussion encompassed morphological characteristics and natural history of sea turtle species relative to environmental conditions and behavioural observations (see Supplementary Material). We used QGIS 3.32 to map the sightings in relation to (i) local bathymetry and (ii) sea surface temperature (SST) for July, 2023 across the North Pacific (see Supplementary Material).

D. Johnston sighted the turtle from a kayak at approximately 10:30 AM near the drop-off of the Monterey Submarine Canyon. The turtle came up to breathe (36.808711°, −121.805436°), swam southward, and was seen approximately five minutes later (36.803556°, −121.804919°) (Figure 2). The turtle was active and amid potential prey species, but not observed eating (Supplementary Table 1). The pelagic location with possible upwelling and high invertebrate abundance corresponds with suitable foraging conditions for loggerheads (Figure 2, Supplementary Table 1). The proportionally large head, tannish-brown coloration, limited number of costal scutes, heart-shaped carapace, and limited number of prefrontal and tympanic scales (Supplementary Table 1) coincide with loggerhead morphology (Lee et al., 2014).

**Figure 2:**
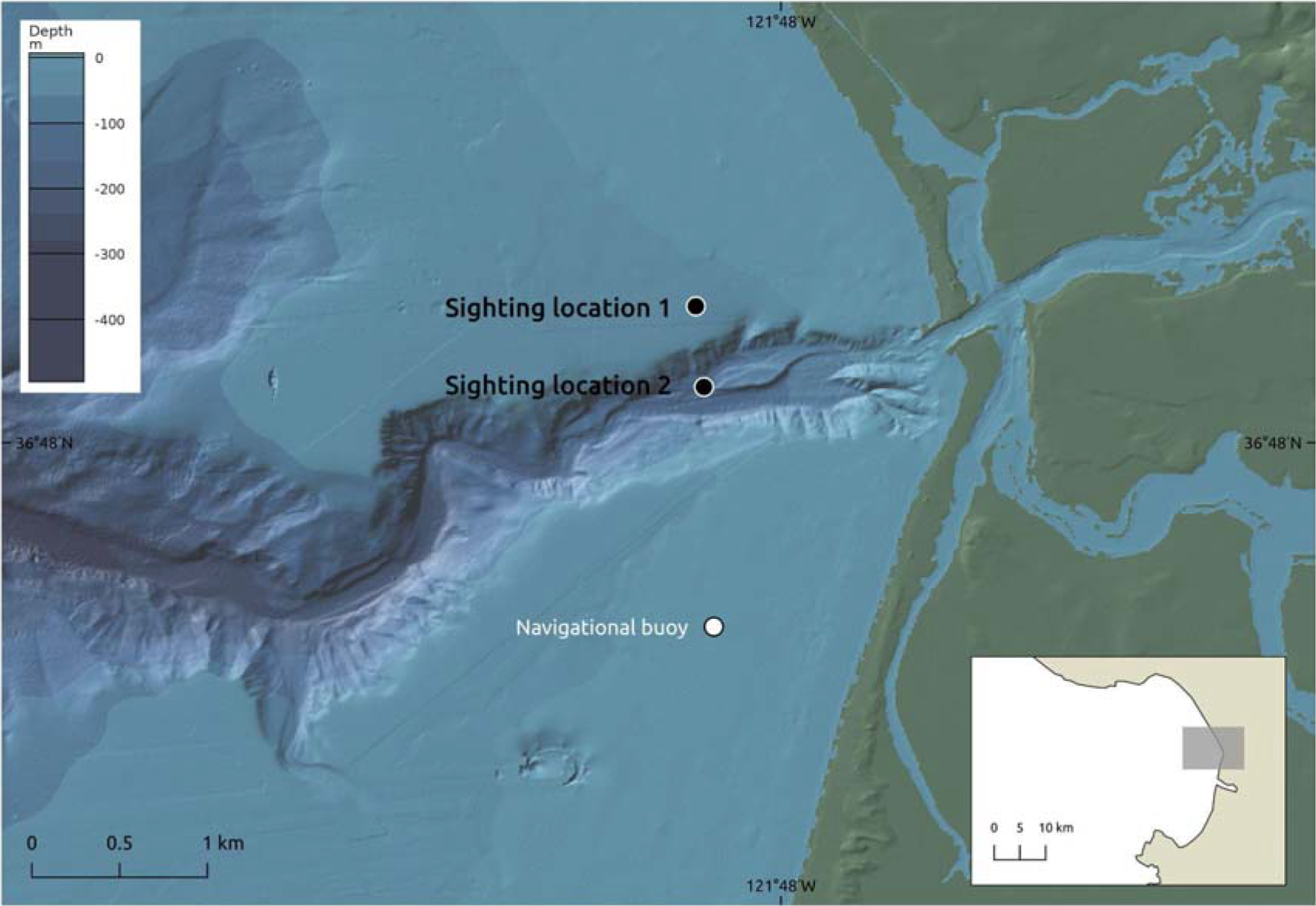
Monterey Bay loggerhead (*Caretta caretta*) sighting locations, with high-resolution (10m) bathymetry of Monterey Submarine Canyon (Paull & Caress, 2019). Figure made with GeoMapApp 3.7.1, CC-BY (Ryan et al., 2009).

Local SST during the sighting (12.5 °C) (NOAA, 2023b) was within the low range of reported loggerhead thermal tolerance levels, which range from 10-27.84 °C, with likely cold-stunning at <8 °C (Briscoe et al., 2016). Prolonged loggerhead presence has been recorded at temperatures as low as 10.1°C (Casale et al., 2012). Loggerhead movement is strongly correlated with SST variations, especially ENSO dynamics which may drive transient loggerhead presence in the temperate North Pacific (Figure 1). Loggerhead strandings have been reported in British Columbia, Canada, and the Gulf of Alaska during El Niño events (Halpin et al., 2018), and 15,000+ loggerheads were recorded in the Southern California Bight (SCB) during the 2015 El Niño event (Eguchi et al., 2018; Briscoe et al., 2021) (Figure 1). Loggerheads’ close association with SST has led to dynamic-fisheries management policies for by-catch reduction in the SCB through a time-area closure during El Niño events (Eguchi et al., 2018). At the time of the sighting, there was an active El Niño advisory for December 2023-February 2024 (NOAA, 2023c).

This finding highlights the pivotal role of LEK and CS in documenting species occurrences beyond their anticipated habitats during climatic anomalies, and the importance of robust methods for recording and identifying species when photographic or video evidence is lacking. Importantly, predicting distribution shifts in rare species using only conventional sampling methods may spatially bias predictions and prove problematic in rapidly changing conditions (Taheri et al., 2021; Jeliazkov et al., 2022). LEK and CS can be reliably integrated into scientific research using established methods from ethnobiology to assure quality, clarity, and reliability, and generate high-quality data with broad coverage for rare or elusive species (Albuquerque et al., 2014; McKinley et al., 2017). Furthermore, LEK and CS can inform dynamic management under climate change, while integrating diverse voices into conservation science.

## Author contributions

Study design, fieldwork: MMEC, GGM, DJ; data analysis: MMEC; figures: MMEC; writing or reviewing manuscript: MMEC, NC, JS, LC, GGM; final approval: all authors.

## Acknowledgements

We thank the Society for Conservation Biology for funding through the David H. Smith Conservation Research Fellowship awarded to MMEC.

## Ethical standards

Fieldwork followed the International Society of Ethnobiology (ISE) Code of Ethics (2008).

## Data availability

Primary data are encrypted and only accessible to the research team per ISE guidelines.

