## Supplementary Material for "Local Ecological Knowledge enhances our capacity to document potential climate sentinels: a loggerhead sea turtle (*Caretta caretta*) case study"

DAVID JOHNSTON Kayak Santa Cruz, Santa Cruz, CA, U.S.A.

BIORXIV PREPRINT

### HARD-SHELLED TURTLE SIGHTINGS

Other recorded hard-shelled sea turtle sightings in Monterey Bay include one live green turtle (*Chelonia mydas*) in 2017 (NOAA, unpublished data); a live, stranded olive ridley (*Lepidochelys olivacea*) in 2011 (Monterey Bay Aquarium, 2011); and live olive ridleys in 1969 (Reichart, 1993) and August, 2023 (The Marine Mammal Center, 2023).

### METHODS

The presumptive loggerhead sighting was reported by co-author D. Johnston, a seasoned naturalist with 40 years of experience and ~16,000 hours of observation time in Monterey Bay. M.M. Early-Capistrán conducted a telephone interview on the day of the sighting, and on July 13 M.M. Early-Capistrán and G. Garibay-Melo conducted a semi-structured interview (Bernard, 2011; Albuquerque et al., 2014). We also employed participatory modelling, which uses mapping exercises to capture and co-produce spatial representations of LEK (Gray et al., 2017; Wedemeyer-Strombel et al., 2019). Interviews were carried out following ethical protocols of the International Society of Ethnobiology (2008). The interview and participatory modelling were audio- and video-recorded and transcribed with informed consent, and coordinates from collaborative maps were recorded in Google Earth Pro 7.3.6.

We followed established ethnobiological techniques for species identification via visual stimulus (Albuquerque et al., 2014). We reviewed three sea turtle identification guides, one of which was designed for community scientists observing sea turtles from above-water (e.g., cruise ships, tour boats, and other vessels) (Pritchard & Mortimer, 1999; Secretariat of the Pacific Community, 2003; Upwell & Grupo Tortuguero de las Californias, 2023), and discussed (i) which species had morphological characteristics most similar to the observed turtle and (ii) the habitat preferences and natural history of each species in relation to the environmental conditions and behavioural observations related to the sighting (e.g., where each species is habitually found, dietary preferences, migratory patterns, etc.). We reviewed composites of photos of the three potential species (*L. olivacea, C. mydas*, and *C. caretta*). Leatherback sea turtles (*Dermochelys coriacea*) also forage in the Monterey Bay region, but were ruled out due to their distinctive morphology (Benson et al., 2020). Each composite image contained photographs of a single species from multiple angles, and species names were not provided during the discussion to avoid biassing responses. In-depth discussion included factors such as the size and number of scutes, facial scaling, pigmentation, cranial morphology, and the size and proportions of the axial body and appendages.

We transferred coordinates from participatory modelling to GeoMapApp 3.7.1 (Lamont-Doherty Earth Observatory, 2023) to access Global Multi-Resolution Topography synthesis (Ryan et al., 2009) and high-resolution (10m) bathymetric data of Monterey Canyon (Paull & Caress, 2019). Additionally, we used NASA monthly sea surface temperature (SST) products for July, 2023 collected with Moderate Resolution Imaging Spectroradiometer (MODIS) instruments at 0.1° resolution based on MODIS-calibrated mid- and far-infrared (IR) radiances (Bands 20, 22, 23, 31, and 32 from MOD02) to map the sighting location in relation to frequently used foraging habitats (NASA Earth Observatory, 2023). All maps were generated with QGIS 3.32.

BIORXIV PREPRINT

SUPPLEMENTARY TABLE 1 Key observations of presumptive loggerhead turtle (*Caretta caretta*) sightings from Local Ecological Knowledge (semi-structured interview and participatory modelling).

| | Quote (D. Johnston) |
|---|---|
| **Oceanographic features** | |
| | "It was straight off of Moss Landing. It was probably right over where the submarine canyon begins and so it could have been a couple hundred feet deep or it could have been near the trench and only, you know, 60 or 80 feet deep, but somewhere around that, where the submarine canyon initiates about a mile from shore [...] If it went over the trench, it could have been a couple hundred feet just for a few hundred yards right there because it's pretty narrow. There could have been some upwelling right there." |
| **Oceanographic conditions** | |
| | "The swell was low, it was less than two feet mixed swell, sunny, not very much wind [...] It was around 10:30 or 11 am [...] [water temperature was] upper 50s, 55 to 60. [...] warmer than usual." |
| **Associated fauna** | |
| | "A lot of pieces of jellyfish around. There haven't been, like, huge swarms of jellyfish yet right there this year, but there was definitely some jellyfish pieces around. Purple stripe jellyfish, brown sea nettles, moon jellies. We did see a big salp, too [...] We saw a mola-mola right then as well. Harbour porpoises. And seals and sea lions and otters." |
| **Morphology** | |
| | "The head appeared to be tan coloured. It didn't have a lot of prefrontal scales that I noticed."<br>"Then I saw the carapace which seemed to have not very many scutes."<br>"The flippers seemed to be pretty large and the shell appeared to be asymmetrical."<br>"[It] was my impression that [the carapace] was more heart-shaped."<br>"Its head [...] was maybe the size of a small harbour seal head, and round."<br>"I could just see, like, the front of the face and it didn't seem like a lot of scales. It just seemed, like, more, like, kind of tan and large scales, two, three scales or something, you know, two on each side." |
| **Pigmentation** | |
| | "The colour of the shell was kind of greyish, tannish brown."<br>"[Its head] was a tan colour and large."<br>"It was that colour [signalling *C. caretta* in identification guide] [...] a tannish brown."<br>"Maybe, like, on the edges of the scales kind of a little reddish." |
| **Size** | |
| | "The carapace was about 30 inches across." |

BIORXIV PREPRINT


BIORXIV PREPRINT
